# Confluence and tight junction dependence of volume regulation in epithelial tissue

**DOI:** 10.1101/2022.03.02.482689

**Authors:** Theresa A. Chmiel, Margaret L. Gardel

## Abstract

Epithelial cell volume regulation is a key component to tissue stability and dynamics. In particular, how cells respond to osmotic stresses is of significant physiological interest in kidney epithelial tissue. For individual mammalian cells, it is well established that Na-K-2Cl cotransporter (NKCC) channels mediate cell volume homeostasis in response to hyperosmotic stress. However, whether mature epithelium respond similarly is not well known. Here we show that while small colonies of MDCK epithelial cells behave similarly to single cells and exhibit volume homeostasis that is dependent on the NKCC channel function, mature epithelial tissue does not. Instead, the cell volume decreases by 33% when confluent monolayers or acini formed from MDCK are subjected to hyperosmotic stress. We show that the tight junction protein, ZO-1, and Rho-associated kinase (ROCK) are essential for osmotic regulation of cell volume in mature epithelium. Since these both are known to be essential for tight junction assembly, this strongly suggest a role for tight junctions in changing volume response in mature epithelium. Thus, tight junctions act either directly or indirectly in osmotic pressure response of epithelial tissue to suppress volume homeostasis common to isolated epithelial cells.

## Introduction

Epithelial cells actively regulate their volume in response to osmotic gradients through management of ion concentration and cytoskeletal tension (Delpire and Gagnon, 2018). Animal cells lack rigid structures that would help to maintain an osmotic gradient across the plasma membrane, meaning that when an osmotic pressure is applied to the membrane, a cell must react to diffuse the osmotic stress on the membrane or risk rupturing the cell membrane(Strange, 1993; Hoffmann *et al*., 2009). It does so primarily by controlling the movement of solutes across the cell membrane through tightly regulated ion channels (Finan and Guilak, 2010). And so, while osmotic stress is a mechanical force on a tissue, how the cells in that tissue respond physically and the regulatory pathways involved in this response is a significantly more complicated relationship.

The pump-leakage model is the basic model that is used to understand regulation of cell volume through ion transport and in response to changes in external or internal osmotic pressure (Strange, 1993). When hypertonic media is introduced, highly membrane permeable water initially rushes out of the cell, causing the cell to shrink in volume. The cell then reacts by activating a variety of transport channels, and predominately the Na-K-2Cl cotransporter (NKCC) family (Haas, 1994), to increase ion concentration in the cytoplasm and restore the osmotic gradient (Haas, 1994; Delpire and Gagnon, 2018). This process to re-establish cell volume after a hyperosmotic shock is known as a regulatory volume increase (RVI).

The response to osmotic gradients in epithelial tissue is not as well understood as in single cells and, presumably, involves both cell and tissue-scale responses. In polarized epithelial tissue, tight junctions assemble at the apical surface and create a barrier to prevent the extracellular flow of osmolytes across the epithelial tissue (Günzel and Yu, 2013; Varadarajan *et al*., 2021). Tight junctions prevent “leakiness” in epithelial tissue and are designed to act as barrier between the internal and external cellular environments (Fischbarg, 2010; Tokuda and Yu, 2019). The integrity of the tight junction is critical for proper organ function (Lee *et al*., 2006, 2018; Liu *et al*., 2012), and the depletion of the tight junction protein zonula occludins-1 (ZO-1) (Odenwald *et al*., 2018) or the cytoskeletal regulator Rho-associated protein kinase (ROCK) (Walsh *et al*., 2001) both result in increased tissue permeability. Renal epithelium in particular are consistently exposed to changing osmotic conditions as solute concentration passing through the kidney is constantly in flux (Beck *et al*., 1998).

Because the tight junctions regulate osmotic flow across the epithelium, they control the special regulation of osmotic stress on the tissue. In addition, polarized renal epithelial tissue confine NKCC1, a member of the NKCC family of ion channels, to the basolateral membrane(Carmosino *et al*., 2008; Koumangoye and Delpire, 2021) and away from any osmotic stresses at the apical surface of the tissue. This functions to create an osmotic pressure gradient not just across a cell’s plasma membrane as we observe in single cells, but a differential pressure gradient across the epithelial tissue, the regulation and cellular response to which is not well understood.

Here, we examine the mechanics of volume regulation in renal epithelial tissue formed from madin darby canine kidney (MDCK) cells and find both a confluence and a tight junction dependence on the tissue’s ability to maintain volume homeostasis under hypertonic conditions. We find that small colonies of epithelial tissue exhibit volume homeostasis in response to hyperosmotic stress, in mechanism reliant on NKCC channels. However, the volume of cells within mature monolayers and acini is acutely suppressed hours after hypertonic conditions were introduced, indicating an inhibition of the process of regulatory volume increase. Disruption of tight junctions in mature monolayers, either through the depletion of ZO-1 or the inhibition of ROCK recover volume homeostasis in response to hyperosmotic stress. Based on these findings, we report a role of the tight junctions in qualitatively modifying cell volume regulation in epithelial tissue.

## Results

### Cell volume in epithelial colonies recovers from hyperosmotic shock via NKCC-mediated regulatory volume increase

To measure the volume response of renal epithelium to osmotic stress, we plated MDCK-II cells sparsely on glass and allowed them to grow for 24 hours, creating small colonies of 4-37 cells and imaged them fluorescently with the addition of CellMask Orange membrane stain. We exposed these colonies to either isotonic (normal DMEM media) or hypertonic (an added 200μM of the synthetic sugar sorbitol) conditions for 3 hours examined whether the colony was able to recover its isotonic volume (Figure 1A), i.e. whether the colony was able to undergo regulatory volume increase (RVI) and maintain volume homeostasis. We observed that the average colony cell volume after a long-term hypertonic shock is slightly but not significantly reduced when compared to the average volume of colonies in isotonic conditions (Figure 1B). In addition, when we examine a colony immediately after the addition of hyperosmotic media, we see that there is an initial decrease in the average cell volume, followed by almost complete volume recovery consistent with RVI (Figure 1C).

**Figure 1:**
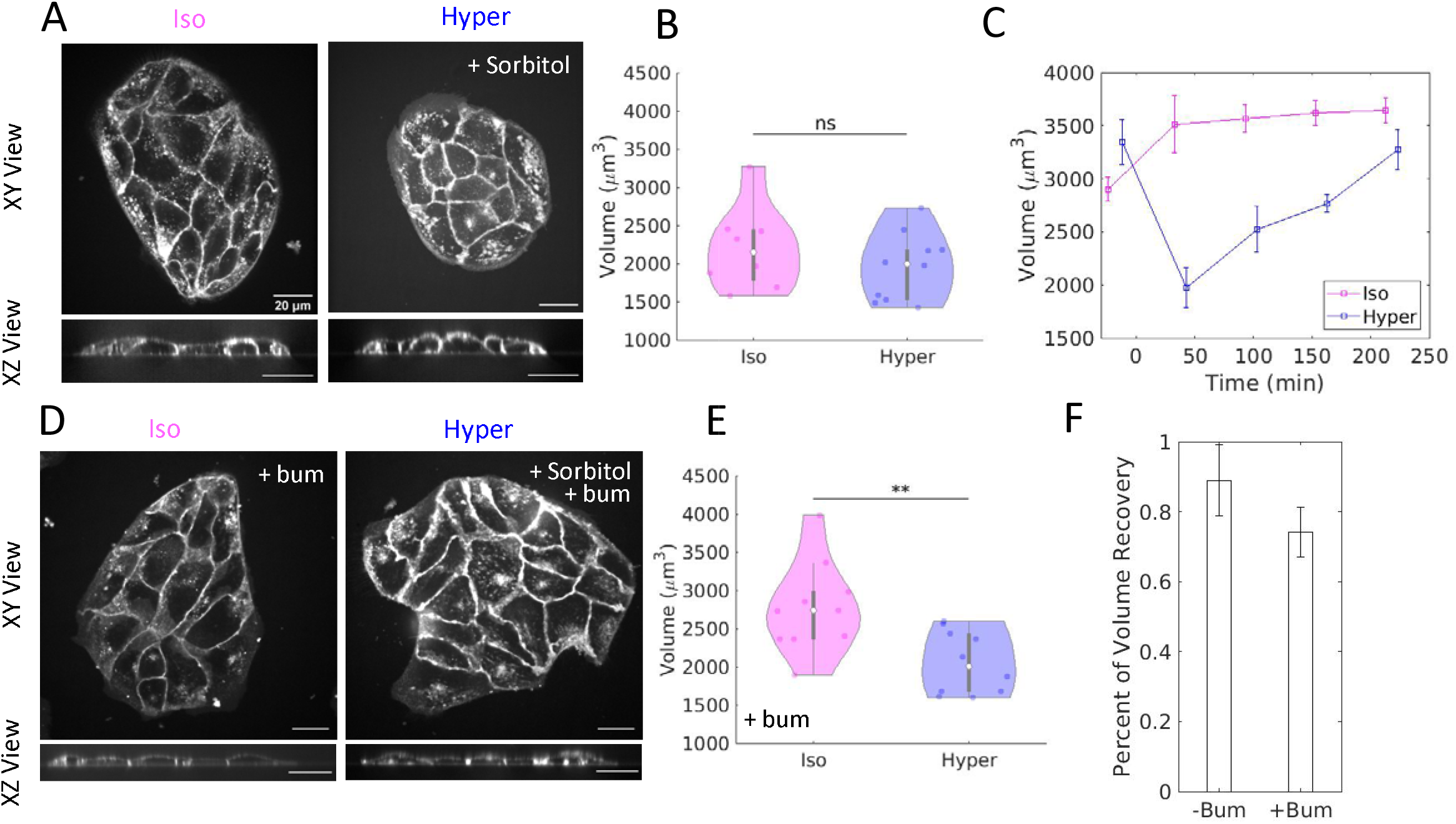
Cell volume in epithelial colonies recovers from osmotic shock via NKCC-mediated regulatory volume increase. (A) Images of live MDCK-II colonies three hours after an osmotic shock in both a top down (XY) view and a side view (XZ). Cell membrane stained with CellMask Orange. Colonies in isotonic (Iso) conditions are in control media, while colonies in hypertonic (Hyper) conditions have 200μM sorbitol added. Scalebars are 20μm. (B) Violin plot of average colony cell volume in isotonic (n=8 colonies) and hypertonic (n=10 colonies) conditions measured three hours after media exchange. Colonies are 4-37 cells in size. ns=p>0.05 as calculated by the student’s t-test. (C) Average colony cell volume in isotonic (Iso) and hypertonic (Hyper) conditions imaged every hour for four hours. Media exchange occurs at t=0min. Error bars represent standard error of the mean. (D) Images of live MDCK colonies three hours after an osmotic shock and the addition of 10μM bumetanide. Cell membrane stained with CellMask Orange. Colonies in isotonic (Iso) conditions are in control media, while colonies in hypertonic (Hyper) conditions have 200μM sorbitol added. Scalebars are 20μm. (E) Violin plot of average colony cell volume in isotonic (Iso) and hypertonic (Hyper) conditions measured three hours after media exchange and the addition of 10μM bumetanide. n=10 colonies each with 5-37 cells. **=p<0.01 as calculated by the student’s t-test. (F) Percent of volume recovery for colonies with and without the addition of 10μM bumetanide. Volume recovery is measured by the ratio of mean hypertonic volume to mean isotonic volume three hours following media exchange. Error bars represent standard error of the mean.

We next wanted to confirm that the volume recovery observed is due to ion channel activity as expected for RVI. MDCK cells express exclusively the Na^+^-K^+^-Cl^−^ cotransporter 1 (NKCC1), which has been found to localize basolaterally in epithelial tissue (Mykoniatis *et al*., 2010; Koumangoye *et al*., 2018). We treated colonies with 10μM of bumetanide, a Na^+^-K^+^-Cl^−^ cotransporter (NKCC) inhibitor and saw that volume homeostasis in response to hypertonic conditions is abrogated (Figure 1D). Cell volume is significantly reduced after 3 hours of hypertonic conditions in the presence of NKCC inhibitor, compared to cells in isotonic conditions (Figure 1E). After 3 hours, NKCC inhibition under hyperosmotic conditions resulted in a reduction in average colony cell volume to 74% of isotonic cell volume, while control cells were able to recover to 89% of their initial volume (Figure 1F). This indicates that small colonies of MDCK cells exhibit volume homeostasis mediated by NKCC channels, consistent with well-established regulatory volume increase mechanism that maintains volume homeostasis in single cells in response to hyperosmotic stress.

### Characterization of monolayer volume and volume variation

To explore changes in cell volume occurring in mature epithelium, we develop protocols to characterize the changes in cell volume. MDCK-II cells are plated densely on glass slides and grown for 48 hours with 100% confluency achieved after 24 hours, creating a mature epithelial tissue. CellMask Orange membrane dye is added prior to imaging to clearly outline the apical and basal membranes in monolayers (Figure 2A). This allows us to calculate average monolayer height for a desired field of view in a sample through plotting the average intensity for each slice of our 3-dimensional image and fitting the two peaks in intensity, which represent the apical and basal membranes of the sample, to a gaussian (Figure 2B). The difference between these two peaks is the local average height of the monolayer. We then measure the average cross-sectional area and approximate average cell volume in a monolayer as the average cross-sectional area multiplied by the average monolayer height (Figure 2C).

**Figure 2:**
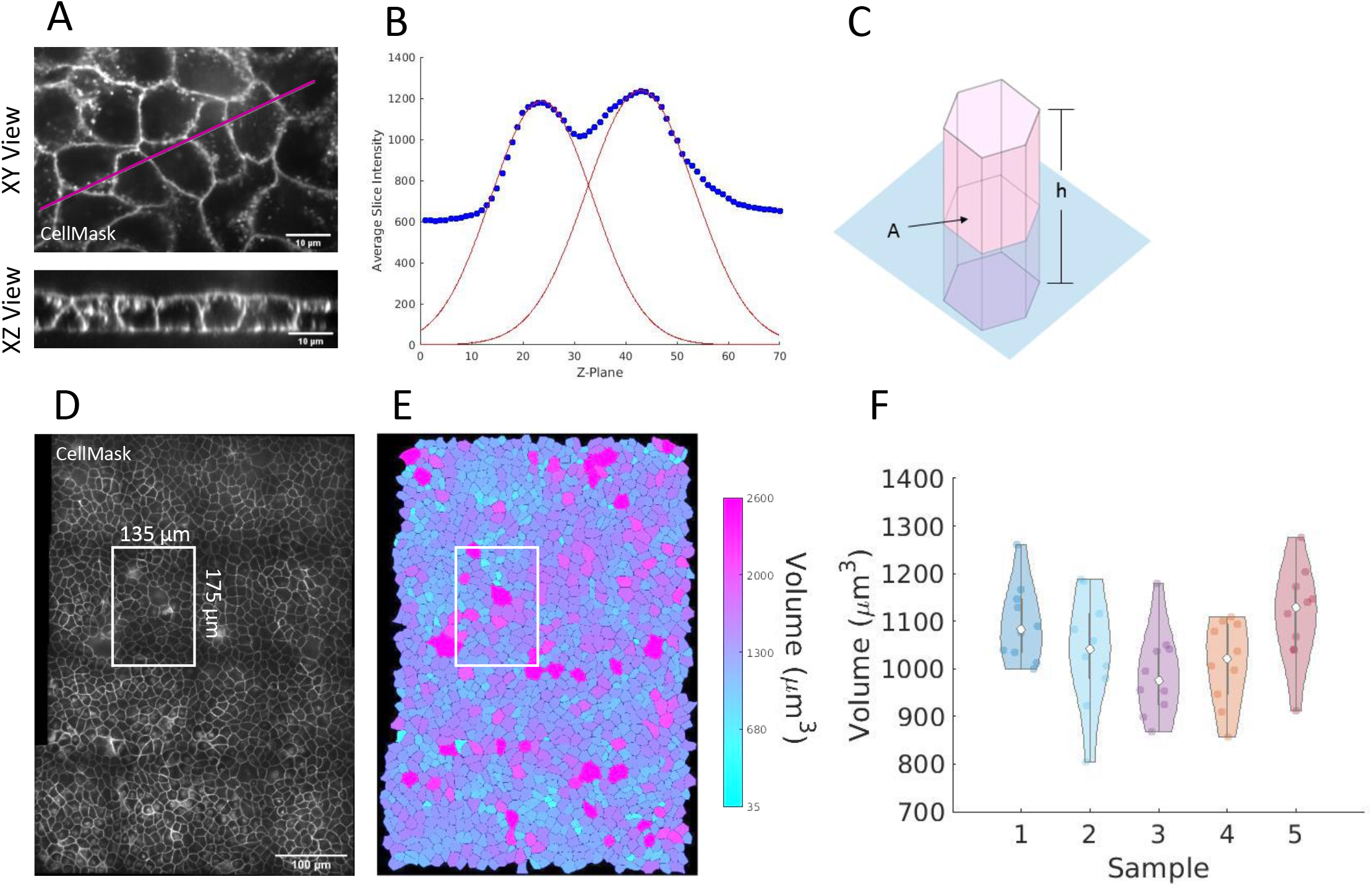
Characterizing Monolayer volume and volume variation. (A) Images of live MDCK monolayers. Cell membrane stained with CellMask Orange. Scalebars are 10μm. (B) Intensity of CellMask Orange membrane stain at different slices of the three-dimensional image, used to measure average monolayer cell height. Average monolayer volume is approximated as average height, h, multiplied by average cross-sectional area, A. (D) A widefield view of a monolayer of MDCK cells compared to the 135×175μm^2^ field of view used as a sample unit when measuring local volume in a monolayer. (E) The volume of each individual cells in the widefield monolayer. (F) Violin plot representing the variation in volume across monolayers. Each sample represents a difference monolayer with each data point representing a different 135×175μm^2^ field of view in that monolayer.

MDCK monolayers exhibit natural variation in both cell density and cell volume (Zehnder *et al*., 2015) (Figure 2D) and the scale of this variation can be seen within a single 135×175μm^2^ field of view (Figure 2E). We chose this range to average over because the natural variation in local volume is consistent with the average variation in volume between one monolayer and another (Figure 2F), allowing us to treat each field of view’s local volume as independent.

### Mature epithelial tissue does not recover from a long-term osmotic shock

We next perform iso and hyper-tonic experiments on mature epithelium formed by plating MDCK cells at high density onto glass and incubating for 48 hours. Under these conditions, tight junctions form in this polarized model epithelial tissue to facilitate well-characterized barrier function (Lee *et al*., 2006; Fischbarg, 2010). In this condition, the hyperosmotic media presumably remains confined to the apical cell surface. Upon media exchange to hypertonic conditions, we discover that that average volume of the cells decreases by 20-35% and does not recover over time such that the epithelial height and volume remain permanently reduced even after three hours (Figure 3, A-C).

**Figure 3:**
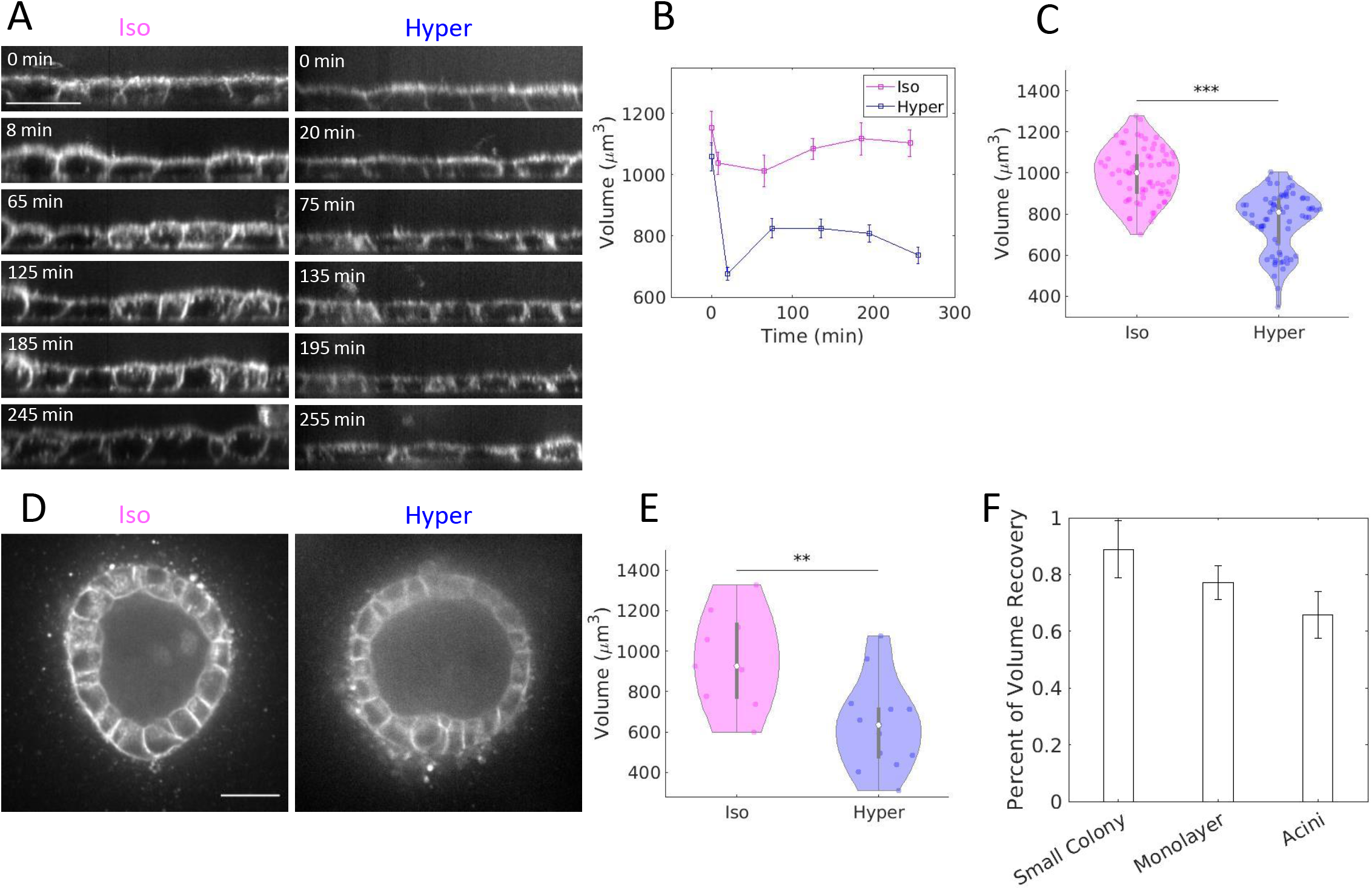
Mature epithelial tissue does not recover from a long-term osmotic shock. (A) Images of live XZ views of MDCK monolayers taken over five hours. Cell membrane stained with CellMask Orange. Monolayers in isotonic (Iso) conditions are in control media, while those in hypertonic (Hyper) conditions have 200μM sorbitol added. Scalebars are 20μm. (B) Average cell volume in isotonic (Iso) and hypertonic (Hyper) conditions imaged every hour for four hours. Media exchange occurs at t=0 min. Error bars represent standard error of the mean. n=10 fields of view from one monolayer. (C) Violin plot of average monolayer cell volume in isotonic and hypertonic conditions measured three hours after media exchange. n=70 fields of view across 7 separate monolayers of varying density with 100-165 cells per field of view. ***=p<0.0001 as calculated by the student’s t-test. (D) Cross sections of hollow MDCK acini grown in collagen gel. Cell membrane stained with CellMask Orange. Acini in isotonic (Iso) conditions are in control media, while those in hypertonic (Hyper) conditions have 200μM sorbitol added. Scalebars are 20μm. (E) Violin plot of average acini cell volume in isotonic (n=9 acini) and hypertonic (n=13 acini) conditions measured three hours after media exchange. **=p<0.01 as calculated by the student’s t-test. (F) Volume recovery is measured by the ratio of mean hypertonic volume to mean isotonic volume three hours following media exchange. Error bars represent standard error of the mean.

To determine whether mature epithelial tissue reacts similarly independent of configuration, we performed these experiments in MDCK acini in collagen gel. MDCK acini are confluent cysts that are formed by seeding cells sparsely within collagen gel and incubating for eight days. Acini are polarized with the apical surface at the inner surface of the cyst known as the lumen and the basal surface facing the external media. Thus, for acini, the basal and basolateral cell surfaces are exposed to the exchanged media and the barrier prevents transport to the apical surface. Similar to confluent monolayers, the hyperosmotic shock permanently reduces volume of cells within acini by 34% (Figure 3, D and E).

Comparing these data, hyperosmotic shock leads to 20-35% cell volume reduction in both confluent monolayers plated on glass and MDCK acini grown in collagen gel. This volume reduction is not seen in MDCK colonies, which recover to 90% of their isotonic volume within three hours (Figure 3F). This allows us to conclude that the NKCC-mediated regulatory volume increase after hyperosmotic shock that preserves volume homeostasis in single cells and small colonies is hampered in confluent renal epithelial tissue.

### Tight junctions are required to prevent volume recovery in mature epithelial tissue

We speculated that the differences in volume regulation between small colonies and mature epithelium might arise from tight junctions. Tight junctions, which are critical for barrier function, are not observed in small colonies and assemble in rho-kinase (ROCK) and ZO-1 mediated processes in confluent and polarized epithelium (Hirase *et al*., 2001; Walsh *et al*., 2001; Odenwald *et al*., 2018). To assess this, we first formed monolayers and acini from MDCK-II cells with the tight junction proteins ZO-1 and ZO-2 knocked-down (Choi *et al*., 2016). Interestingly, we found that that cell volume recovered after hyperosmotic shock in these cells. Similar to that observed for small colonies, the volume of ZO-1/ZO-2 deficient cells in mature epithelium acutely decreased but recovered over the subsequent several hours (Figure 4, A and B). After 3 hours, the average cell volume is not significantly different than those in isotonic media (Figure 4C). Thus, ZO-1 is critical to the differing volume regulation observed in confluent epithelial tissue.

**Figure 4:**
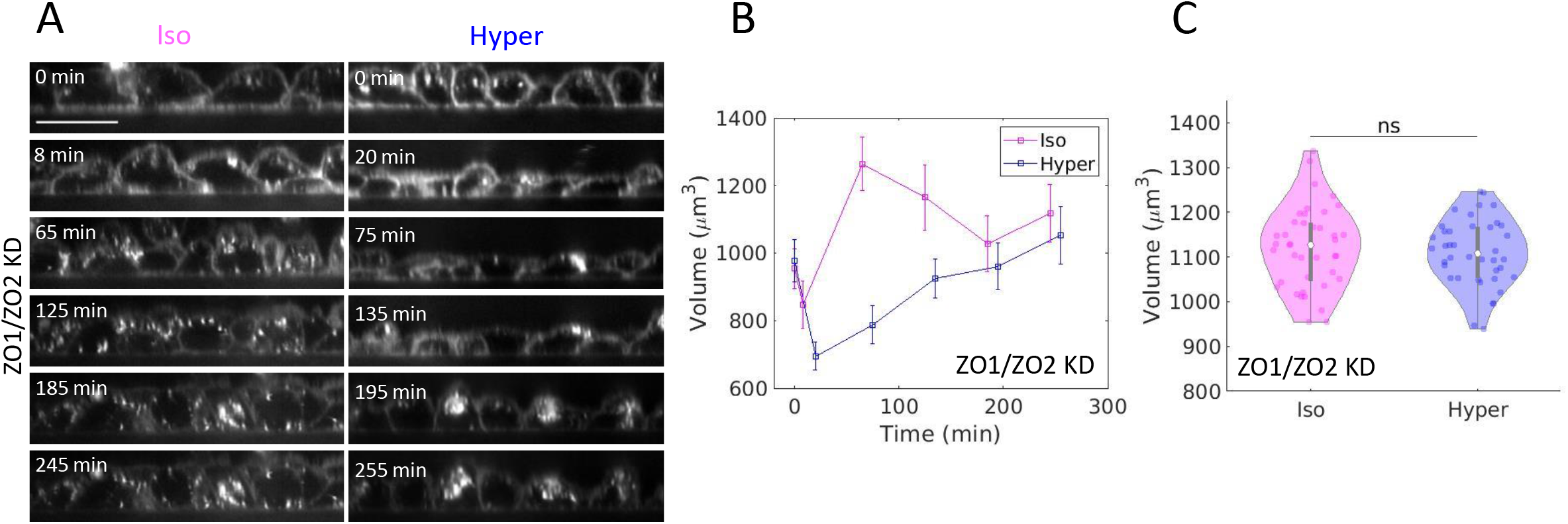
The tight junction protein ZO-1 is required to prevent volume recovery in mature epithelial tissue. (A) Images of live XZ views of ZO-1/ZO-1 knock-down (KD) MDCK monolayers taken over five hours. Cell membrane stained with CellMask Orange. Monolayers in isotonic (Iso) conditions are in control media, while those in hypertonic (Hyper) conditions have 200μM sorbitol added. Scalebars are 20μm. (B) Average cell volume of ZO-1/ZO-1 KD monolayers in isotonic (Iso) and hypertonic (Hyper) conditions imaged every hour for five hours. Media exchange occurs at t=0 min. Error bars represent standard error of the mean. n=10 fields of view in one monolayer. (C) Violin plot of average ZO-1/ZO-1 KD monolayer cell volume in isotonic and hypertonic conditions measured three hours after media exchange. n=40 fields of view across 4 separate monolayers of varying density with 99-150 cells per field of view. ns=p>0.05 as calculated by the student’s t-test.

Another means to alter tight junction assembly is through inhibition of ROCK activity (Hirase *et al*., 2001; Walsh *et al*., 2001) with the addition of 25μM of the ROCK inhibitor Y-27632 during monolayer formation. We find that ROCK inhibition also recovered the process of regulatory volume increase of cells within confluent monolayers or acini that are subjected to hyperosmotic shock (Figure 5, A-E) Taken together, these data show that ROCK inhibition facilitates an 88% volume recovery in monolayers, and 94% for acini (Figure 5F). And so, we are able to conclude that functional tight junctions are critical for the different response to hyperosmotic shock in mature epithelium than in isolated cells.

**Figure 5:**
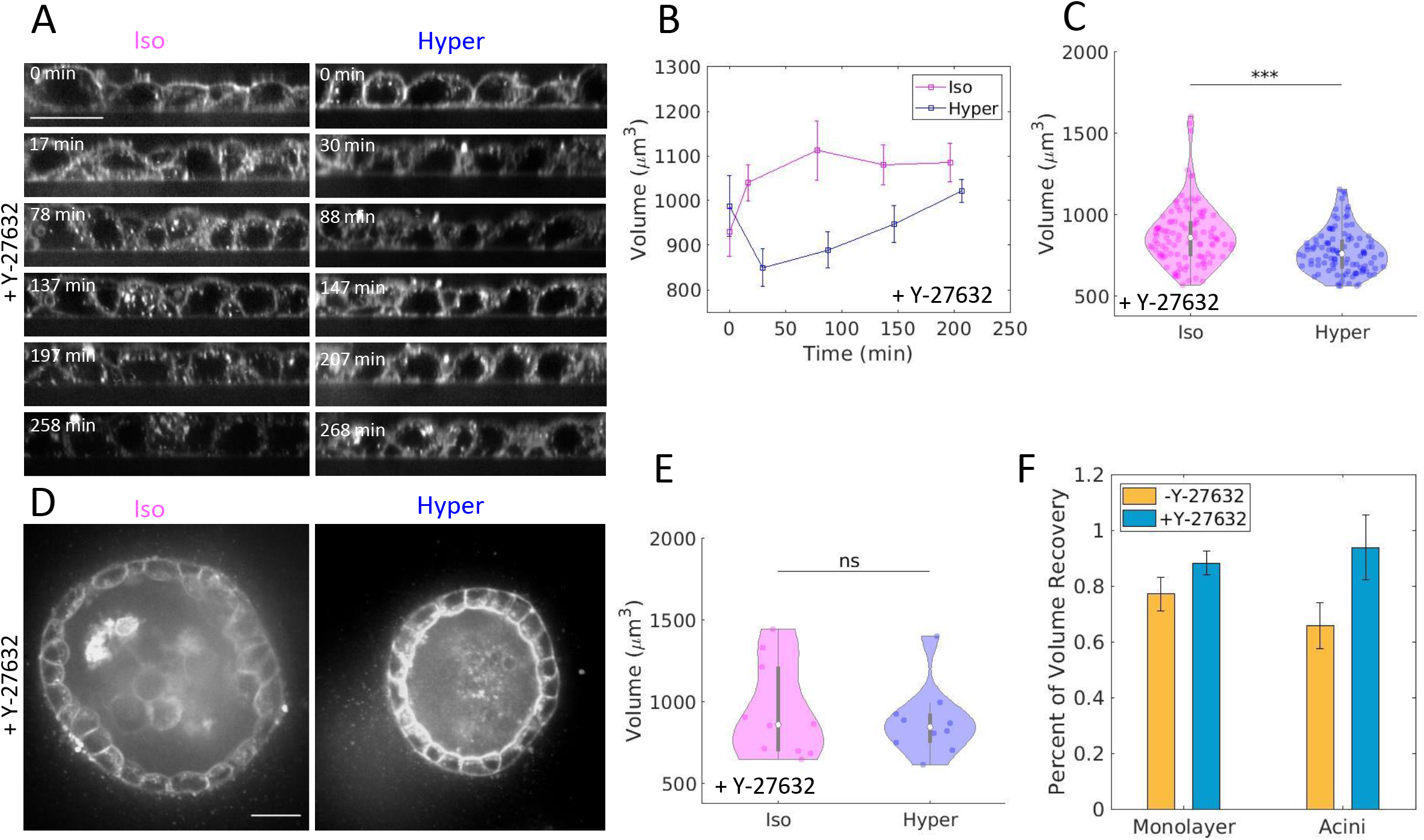
ROCK is required to prevent volume recovery in mature epithelial tissue. (A) Images of live XZ views of MDCK monolayers with 25μM of the ROCK inhibitor Y-27632 taken over four hours. Cell membrane stained with CellMask Orange. Monolayers in isotonic (Iso) conditions are in control media, while those in hypertonic (Hyper) conditions have 200μM sorbitol added. Scalebar is 20μm. (B) Average cell volume of monolayers with 25μM Y-27632 in isotonic and hypertonic conditions imaged every hour for four hours. Media exchange occurs at t=0min. Error bars represent standard error of the mean. n=10 fields of view in one monolayer. (C) Violin plot of average monolayer cell volume with 25μM Y-27632 in isotonic and hypertonic conditions measured three hours after media exchange. n=110 fields of view across 11 separate monolayers of varying density with 40-160 cells per field of view. ***=p<0.0001 as calculated by the student’s t-test. (D) Cross sections of hollow MDCK acini grown in collagen gel. with 25μM Y-27632 added three hours prior to imaging. Cell membrane stained with CellMask Orange. Acini in isotonic (Iso) conditions are in control media, while those in hypertonic (Hyper) conditions have 200μM sorbitol added. Scalebars are 20μm. (E) Violin plot of average acini cell volume in isotonic (n=10 acini) and hypertonic (n=10 acini) conditions measured three hours after media exchange. ns=p>0.05 as calculated by the student’s t-test. (F) Volume recovery is measured by the ratio of mean hypertonic volume to mean isotonic volume three hours following media exchange. Error bars represent standard error of the mean.

## Discussion

The mechanics regulating cell volume homeostasis are vital to the proper functioning of epithelial tissue in animals. Here we find that small colonies of renal epithelial tissue maintain their volume homeostasis and undergo NKCC dependent regulatory volume increase in a manner consistent with the pump-leakage model found in many types of individual cells. Here we find this is not the case for mature epithelium. Following a long-term hyperosmotic shock, the volume of individual cells within mature epithelium is permanently decreased (Figure 6A). Thus, regulatory volume increase that allows for cell volume homeostasis in response to hyperosmotic stress does not occur in mature monolayers. This lack of RVI is dependent on the tight junction assembly. And so, we report an important role of the tight junctions in assisting cell volume regulation in epithelial tissue.

**Figure 6:**
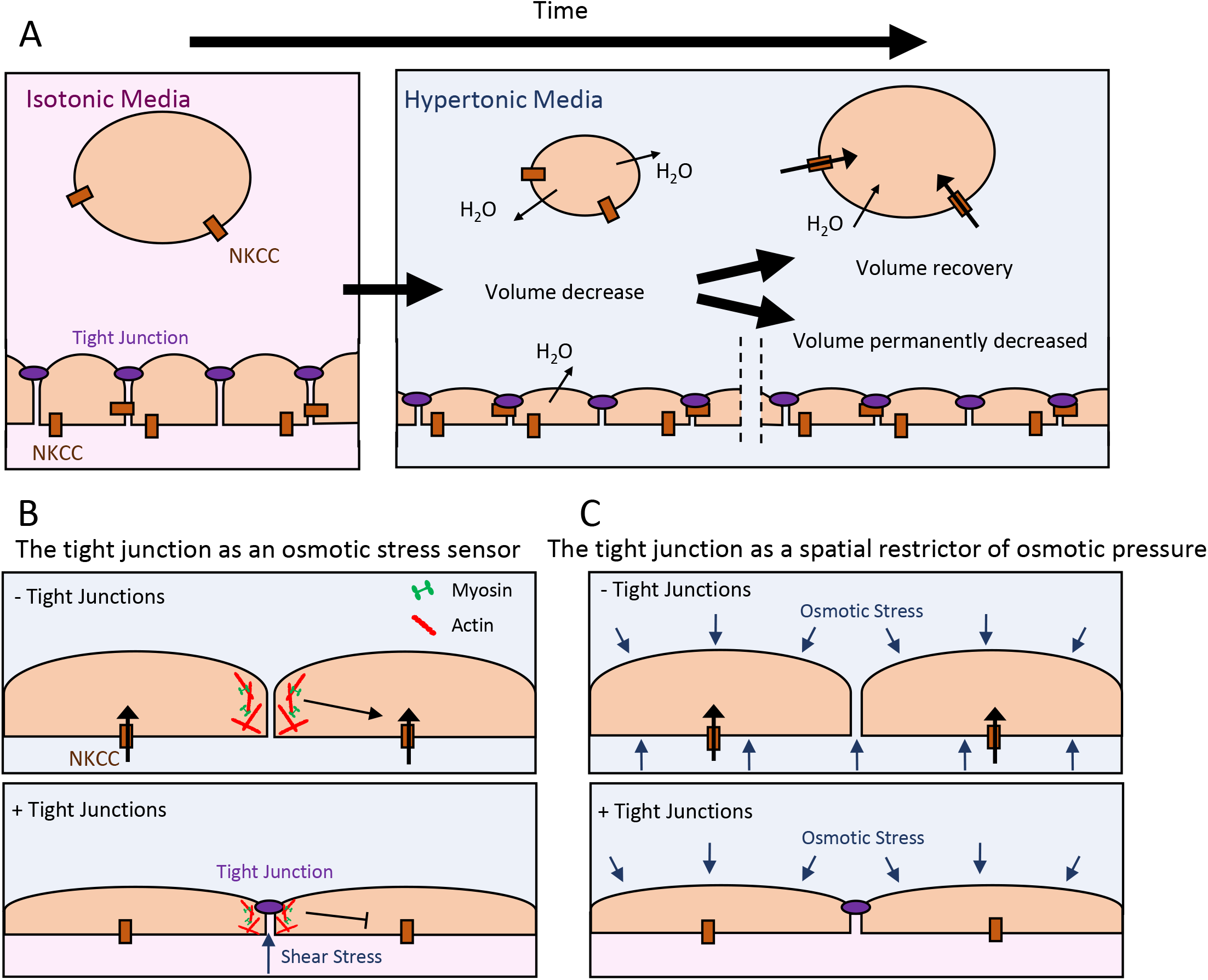
Tight junction regulation of volume homeostasis. (A) A schematic depicting the dynamics of volume homeostasis in single cells and in mature epithelial tissue. With the addition of hyperosmotic media, single cells first shrink in volume as water exits the cell to the surrounding media. Over time, NKCC ion channels pump ions back into the cells, allowing water to follow the osmotic gradient and restore the cell’s volume. A mature epithelium exposed to hypertonic media also reduces in volume as water exits into the surrounding media. However, the NKCC ion pumps are inactive and no RVI occurs. (B) A schematic of the potential role the tight junction plays as an osmotic stress sensor. In the case of epithelium with permeable, i.e. leaky, tight junctions (upper), the absence of the tight junction allows volume homeostasis to occur through NKCC ion channels. In confluent tissue with functioning tight junctions (lower) the sheer stress across the tight junction is transmitted to the mechanosensitive actomyosin cytoskeleton. This results in the inhibition of volume homeostasis, and cell volume does not recover. (C)A schematic of the possible role the tight junction plays as a spatial restrictor of osmotic stress. In this model, epithelia with permeable tight junctions (upper) have zero osmotic gradient across the epithelium, and thus are surrounded at all membranes by hyperosmotic stresses, promoting regulatory volume increase reminiscent of single cell volume homeostasis. In contrast, epithelia with impermeable tight junctions (lower) create a barrier preventing hyperosmotic media from reaching across to both sides of the epithelia, creating a nonzero osmotic gradient and spatially restricting where osmotic pressure is applied to the cells in the epithelium.

Here, we surmise two possible explanations for how the tight junctions may be playing this regulatory role in epithelial tissue. One involves the tight junction as an osmotic sensor, able to communicate hydrostatic pressure across the barrier to the cytoskeleton, while the other considers the osmotic barrier created by the tight junction and how it affects osmotic pressure across epithelial tissue.

It has been previously hypothesized that that the tight junction acts directly in mechanotransduction of osmolarity gradients and regulation of cell volume (Tokuda and Yu, 2019). Osmotic stress locally impact shear stress at the tight junction and, via, action on mechanosensitive cytoskeleton may impact the activity of ion channels regulating regulatory volume increase (Figure 6B). Without functioning tight junctions, sub-confluent or inhibited tissue may lose its ability to detect the osmotic pressure gradient due to the hypertonic conditions, resulting in the downstream effect of active NKCC ion channels facilitating RVI. In contrast, fully confluent and uninhibited epithelial monolayers sense this shear stress at their “leak-proof” tight junctions, and this results in the suppression of volume homeostasis.

An equally interesting explanation is that an intact tight junction serves to confine the osmotic pressure to only a small subsection of the epithelial membrane (Figure 6C). Meaning that instead of an epithelial cell being surrounded by hyperosmotic pressure at all membranes as we see when examining single cell volume regulation, the tight junction restricts that osmotic pressure to only one side of the epithelial tissue, and impact regulatory volume increase. In our results, mature monolayers with functional tight junctions are only exposed to hyperosmotic pressure at the apical membrane and acini are only exposed at the basolateral membrane. Thus, our data does not indicate that the geometry of the osmotic shock is important for this response, as one might expect given known localization of the NKCC1 channel to the basolateral surface (Mykoniatis *et al*., 2010; Koumangoye *et al*., 2018). Instead, it suggests cells react differently to osmotic pressure gradients across the apical-basal plane than when to those across the cell membrane.

Future work is required to understand the mechanisms controlling of volume regulation in mature epithelium, including how tight junctions impact NKCC ion channel activity. We can, however, conclude that tight junctions in mature epithelium suppress volume homeostasis and, as such, cell volume regulation in epithelial tissue is qualitatively different from that of single cells. Given the known consequences of cell volume regulation for their physiology, these results have significant consequences for control of mechanotransduction pathways in epithelial tissue.

## Materials and Methods

### Cell culture

MDCK-II cells were cultured in Dulbecco’s Modified Eagle’s Medium (DMEM) and supplemented with 10% fetal bovine serum (FBS) (ThermoFisher Scientific), 2mM L-glutamine (Invitrogen), and penicillin-streptomycin (Invitrogen). Cells were incubated in a humidified environment at 37C and 5% C0_2_. MDCK ZO-1/ZO-2 KD cells were generously provided by Mark Peifer (University of North Carolina).

### Osmotic shock treatment

Hyperosmotic media was created through the addition of 200 mOsm of sorbitol (Phytotechnology Laboratories) to 290 mOsm DMEM media to make 490 mOsm hypertonic media.

### Sample creation

To create small colonies, MDCK cell were plated sparsely to coat the glass bottom of a 4-well chamber (Ibidi) and incubated for 24 hours. To create a mature monolayer, MDCK cells were plated densely to coat the glass bottom of a 4-well (for timelapse imaging) or 8-well (for non-timelapse imaging) chamber (Ibidi). The cells were then incubated for 48 hours with a change of media at 24 hours. Acini were formed by sparsely plating MDCK cells in 2mg/mL collagen gel for 8 days with the addition of serum starve (1% FBS) DMEM changed every 48 hours. Inhibitors and osmotic treatments were added 3 hours prior to imaging. ROCK inhibited cells were treated with 25μM Y-27632 (Sigma) 3 hours before live imaging. NKCC inhibited cells were treated with 10μM Bumetanide (Sigma) 3 hours before live imaging.

### Microscopy and live cell imaging

2μl/ml of CellMask Orange (Invitrogen) was added to samples 30 minutes prior to imaging to stain the cell membrane. Samples were imaged on an inverted T-E microscope (Nikon) with a confocal CSU-X spinning disk (Yokogawa Electric Corporation), a stage controller (Prior), and a CMOS camera (Zyla-Andor). Metamorph software was used to control the microscope and collect images. A stage incubator (Chamlide and Quorum Technologies) with CO-2, humidity, and temperature control was used for timelapse experiments while a stage heater (Nevtek, ASI 400) was used for non-timelapse experiments. A 561 nm laser (MPB Communications, VFL-P Series) was used to illuminate the CellMask Orange stain. Images were acquired using a 60x Plan Apo NA water immersion objective with a NA of 1.20 and a WD of 0.31-0.28 (Nikon). Three-dimensional images were collected using z-stacks of 0.25μm steps.

### Height and volume analysis

Both ImageJ and Matlab were used for image analysis of monolayer height and volume. Average monolayer height (excluding ZO1/ZO2 KD monolayers) was measured using Matlab by taking the average image intensity for each z-stack of the 3-dimensional image. The peaks in average image intensity at the apical and basal membrane from the CellMask Orange dye were fitted to a gaussian and the distance between these peaks was measured to find the monolayer height. In ImageJ, the Cell Counter tool was used to determine the density of cells in each sample image. This density measurement was used to determine the average cross-sectional area of the cells in the image. The average volume of cells in the monolayer was found by multiplying the average cross-sectional area by the average monolayer height. Volume measurements for ZO-1/ZO-2 KD monolayers, as well as all colonies and acini, were found using an alternate method to that described above. For ZO-1/ZO-2 KD cells and colonies this is because their extended apical domain is not well characterized by the assumption that the apical membrane is flat, and for acini it is because the cells are not grown on a flat surface. In these cases, average cell volume was found by measuring cell junction height, maximum cell height, and basal area for sample cells. This allowed us to model the volume of the cell as a cylinder that is the height of the cell junction length plus a hemisphere on top.

### Violin Plots

The width of each colored region represents volume kernel density which is an estimation of the probability density function of the average cell volume. The white point represents the median cell volume for all colonies measured, and the grey bar represents the interquartile range, meaning the middle 50% of the volume range.

### Quantification and Statistical analysis

Image analysis and quantification was performed in Fiji, Excel, and Matlab. Matlab was used to perform statistical analysis and calculate statistical significance using two-tailed student t-tests where ns=p>0.05, *=p<0.05, **=p<0.01, and ***=p<0.0001.

## Abbreviations

(NKCC): Na-K-2Cl Cotransporter
(RVI): Regulatory Volume Increase
(ZO-1): Zonula Occludins-1
(ROCK): Rho-associated Protein Kinase
(MDCK): Madin Darby Canine Kidney

## Acknowledgements

This work was supported by funding from NIH RO1 GM104032 and ARO MURI W911NF1410403. We are grateful to Mark Peifer (UNC Chapel Hill) for the ZO-1/ZO-2 KD MDCK cell line, and to Jeffrey Matthews, Eric Delpire, and Patrice Bouyer for valuable discussions.

